# The *Polycomb* group protein MEDEA controls cell proliferation and embryonic patterning in *Arabidopsis*

**DOI:** 10.1101/2020.10.13.338707

**Authors:** Sara Simonini, Marian Bemer, Stefano Bencivenga, Valeria Gagliardini, Nuno D. Pires, Bénédicte Desvoyes, Eric van der Graaff, Crisanto Gutierrez, Ueli Grossniklaus

## Abstract

Establishing the body plan of a multicellular organism relies on precisely orchestrated cell divisions coupled with pattern formation. In animals, cell proliferation and embryonic patterning are regulated by *Polycomb* group (PcG) proteins that form various multisubunit complexes (Grossniklaus and Paro, 2014). The evolutionary conserved *Polycomb* Repressive Complex 2 (PRC2) trimethylates histone H3 at lysine 27 (H3K27me3) and comes in different flavors in the model plant *Arabidopsis thaliana* (Förderer et al., 2016; Grossniklaus and Paro, 2014). The histone methyltransferase MEDEA (MEA) is part of the FERTILIZATION INDEPENDENT SEED (FIS)-PRC2 required for seed development^4^. Although embryos derived from *mea* mutant egg cells show morphological abnormalities (Grossniklaus et al., 1998), defects in the development of the placenta-like endosperm are considered the main cause of seed abortion (Kinoshita et al., 1999; Scott et al., 1998), and a role of FIS-PRC2 in embryonic patterning was dismissed (Bouyer et al., 2011; Leroy et al., 2007). Here, we demonstrate that endosperm lacking *MEA* activity sustains normal embryo development and that embryos derived from *mea* mutant eggs abort even in presence of a wild-type endosperm because *MEA* is required for embryonic patterning and cell lineage determination. We show that, similar to PcG proteins in mammals, MEA regulates embryonic growth by repressing the transcription of core cell cycle components. Our work demonstrates that *Arabidopsis* embryogenesis is under epigenetic control of maternally expressed PcG proteins, revealing that PRC2 was independently recruited to control embryonic cell proliferation and patterning in animals and plants.

## INTRODUCTION

A fundamental question in developmental biology is how cells acquire and maintain their identity over time. Most cell types are specified during embryonesis (Stent, 1985) and PcG proteins play an important role in maintaining cell identity by silencing target genes, including pluripotency factors, while their deregulation is associated with cancer (Laugesen et al., 2016; Loubiere et al., 2019). While mutations affecting PRC2 subunits lead to embryo lethality in animals (Faust et al., 1995; O’Carroll et al., 2001; Oktaba et al., 2008; Pasini et al., 2004), plants lacking PRC2 components do not show severe embryonic phenotypes, and most produce viable offspring (Bouyer et al., 2011; Chanvivattana et al., 2004; Kinoshita et al., 2001). An exception are mutants affecting components of FIS-PRC2, i.e. MEA (Grossniklaus et al., 1998), FERTILIZATION INDEPENDENT ENDOSPERM (FIE) (Ohad et al., 1999), FIS2 (Chaudhury et al., 1997), and MULTICOPY SUPPRESSOR OF IRA1 (MSI1) (Köhler et al., 2003a). Seeds inheriting maternal mutant alleles of *FIS* class genes abort due to a failure in endosperm cellularization and embryonic growth arrest, regardless of the paternal genotype. This maternal effect is observed because *MEA* and *FIS2* are regulated by genomic imprinting, leading to parent-of-origin-dependent allelic expression (Jullien et al., 2006; Kinoshita et al., 1999; Vielle-Calzada et al., 1999). This form of epigenetic gene regulation evolved independently in seed plants and mammals, where it plays a prominent role in the placenta and is required for normal embryonic development (Barlow and Bartolomei, 2014; Ferguson-Smith, 2011). Although mutants affecting FIS-PRC2 components produce embryos with increased cell proliferation and disorganized division planes (Grossniklaus et al., 1998; Ohad et al., 1999), embryo abortion was considered to indirectly result from defects in the endosperm(Kinoshita et al., 1999; Scott et al., 1998). Moreover, *fie* homozygous seeds develop normally (Bouyer et al., 2011), and embryo rescue of *mea* seeds produces wild-type looking albeit sterile plants (Grossniklaus et al., 1998), indicating that FIS-PRC2 is not essential for embryonic development (Kiyosue et al., 1999; Leroy et al., 2007). However, in none of the previous studies could the genotype of embryo and endosperm be uncoupled, due to the complexity of plant reproduction. In plants, male and female multicellular, haploid structures (pollen and embryo sac) each produce two gametes that are usually derived from a single meiotic spore and thus genetically identical. When the pollen delivers the two sperm cells to the embryo sac, double fertilization occurs whereby the female gametes, egg and central cell, fuse with one sperm each to develop into embryo and endosperm, respectively.

## RESULTS

### *MEA* is required for embryogenesis in *Arabidopsis*

In addition to expression in the endosperm, *MEA* transcript was detected in embryos up to torpedo stage (Spillane et al., 2007; Vielle-Calzada et al., 1999). Embryos that develop from fertilized *mea* eggs are larger than wild-type and arrest around the heart stage (Grossniklaus et al., 1998). To unravel the embryonic requirement of *MEA*, we aimed to generate seeds with genetically distinct embryo and endosperm. Such seeds can be obtained by two consecutive single fertilization events: the first is achieved with pollen containing a single sperm that fertilizes either the egg or the central cell (Figure 1A, step1). Incomplete double fertilization allows the embryo sac to attract a second pollen tube (Maruyama et al., 2013) (Figure1A, step2). If the second fertilization event involves genetically distinct pollen, the genotype of endosperm and embryo will differ in a fraction of the seeds (Figure 1A, step3). As first pollen donor we used the *kokopelli* (*kpl*) mutant, which produces some pollen with a single sperm (Ron et al., 2010). We performed a first minimal pollination of wild-type and *mea* homozygous plants with *kpl* pollen carrying the GFP reporter gene under control of the *pRPS5a* promoter (*pRPS5a::GFP*, referred to as *kpl+GFP*), which is active in embryo and endosperm (Weijers et al., 2001). In crosses between wild-type and *kpl+GFP* plants, 80.8% of the seeds underwent double fertilization, producing viable seeds (n=631; Figure S1A). This percentage increased to 95.1% upon a second pollination with wild-type pollen (n=645, Figure 1B, Figure S1A). As expected, pollination of *mea* homozygotes with *kpl+GFP* pollen did not rescue seed abortion since the paternal *MEA* allele is inactive (Grossniklaus et al., 1998; Kinoshita et al., 1999; Vielle-Calzada et al., 1999)(n=692; Figure 1B, Figure S1A). As second pollen donor, we used a transgenic line harboring the constructs *pRPS5a::MEA,* allowing paternal expression of *MEA* soon after fertilization, and *pRPS5a::RFP* (referred to as *MEA-rescue+RFP)*, which complemented 44.8% of the *mea* homozygous seeds (n=665, Figure 1B, Figure S1A). In consecutive pollinations with *kpl+GFP* followed by *MEA-rescue+RFP* pollen, 2.8% of *mea* ovules developed into viable seeds (n=1056, Figure 1B, Figure S1A). Given their fluorescence profile, the seeds were classified as follows (n=837; Figure 1C-F): GFP-positive endosperm and embryo (88.6%, Figure 1C); RFP-positive endosperm and embryo (6.1%, Figure 1D); RFP-positive endosperm and GFP-positive embryo (3.1%, Figure 1E); and GFP-positive endosperm and RFP-positive embryo (2.2%, Figure 1F). The latter two classes originated from two single fertilization events resulting in seeds harboring genetically distinct embryo and endosperm as only one of them carried *MEA-rescue*. Eight days after the consecutive pollinations, when wild-type seeds harbored embryos at the bent cotyledon stage (Figure 1G), we observed five phenotypic classes among cleared seeds of *mea* siliques (n=1235; Figure 1H-L): *mea*-looking seeds with embryos arrested around the heart stage and uncellularized endosperm (84.3%, Figure 1H); seeds without visible embryos (5.6%, Figure 1I) originating from either single fertilization events of the central cell or autonomous endosperm development; wild-type looking seeds (4.2%, Figure 1J) derived from double fertilization with *MEA-rescue+RFP* pollen; seeds with abnormal embryos arrested around the heart stage and cellularized, wild-type looking endosperm (2.7%, Figure 1K); and seeds with wild-type looking embryos at the walking-stick stage surrounded by uncellularized endosperm (3.2%, Figure 1L, Figure S1C). Single-seed genotyping of the last class by droplet digital PCR (ddPCR) confirmed that they carried both the GFP and RFP transgenes (n=46; Figure S1D). Given that *kpl+GFP* alone does not rescue defective *mea* embryos, these seeds must have developed from *mea* embryos carrying *MEA-rescue* (RFP-positive) surrounded by *mea* mutant endosperm (GFP-positive).

**Figure. 1.**
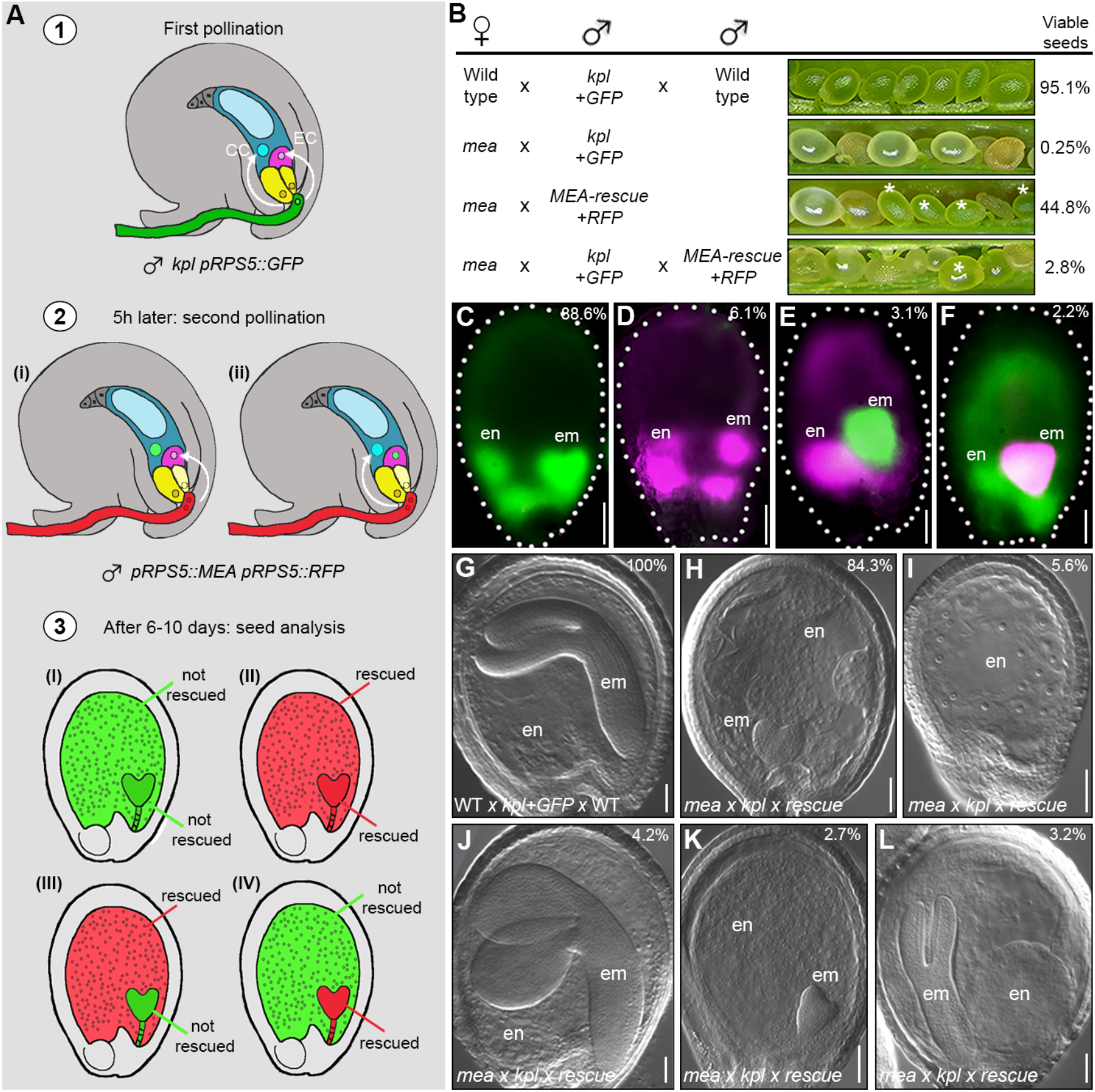
Genetically uncoupling embryo and endosperm development reveals the requirement of *MEA* for embryogenesis. **A.** Schematic representation of the strategy. EC, egg cell; CC, central cell. **B.** Set of crosses performed, with opened siliques showing developing seeds. Asterisks indicate developing seeds among aborting ones. **C-F.** Fluorescent microscopy images of seeds derived from the double pollination experiment: no rescue (C), complete rescue (D), endosperm only rescue (E), and embryo only rescue (F).. **G-L.** Clearing of seeds of the wild-type *x kpl+GFP x* wild-type cross (G) and the five phenotypic classes (H-L) observed in *mea × kpl+GFP x MEA-rescue+RFP* crosses: *mea*-like (H), endosperm only (i), *MEA*-rescued (J), abnormal embryo and wild-type-looking endosperm (K), wild-type-looking embryo and *mea* endosperm. Top right corner: percentage of seeds showing the phenotype. em, embryo; en, endosperm. Scale bar: 50μm

Altogether, these results demonstrate that embryo and endosperm development can be uncoupled in *mea* mutant seeds, and that development of *mea* embryos arrests regardless of the genotype of the endosperm (Figure 1K). Our analysis revealed that the failure in endosperm proliferation and cellularization in *mea* seeds does not cause abortion of the embryo because *MEA*-rescued embryos, even if surrounded by *mea* endosperm, complete development and produce viable progeny (Figure S1C). Thus, *MEA* activity is directly required for normal embryogenesis.

### *mea* embryos display patterning defects, particularly in the root apical meristem

Embryos originating from *mea* eggs (referred to as *mea* embryos) develop as disorganized mass of cells with small, asymmetric cotyledons and an enlarged root meristem (Grossniklaus et al., 1998). To characterize the defects at cellular resolution, we performed a modified pseudo-Schiff Propidium Iodide staining (mPS-PI) of siliques from *mea/MEA* heterozygotes (Figure 2A-C, Figure S2A-B), where a 1:1 segregation of wild-type and *mea* embryos is expected. At the globular stage, 29.2% of the embryos showed ectopic cell divisions in the basal part of the embryo, such that they became almond-shaped (n=48; Figure 2A-B, Figure S1A-B). Disorganized and excessive cell divisions are characteristic for the basal part of *mea* embryos throughout development (Figure 2A-C, Figure S2A-C). At late heart stage, when wild-type embryos contained five to seven cells in the columella and quiescent center (QC) region, this region contained between seven and 18 cells in *mea* embryos (n=120; Figure 2C, Figure S2D). Consequently, the pyramidal organization typical of this part of wild-type root meristems was replaced by a mass of globularly arranged cells in *mea* embryos (Figure 2C, Figure S2B). Consistent with root meristem defects, when grown on vertical plates, *mea* seedlings exhibited severe agravitropic growth and the primary root made loops, upward turns, displayed twisted epidermal cells, and an altered columella root tip organization (n= 76; Figure 2D-I, Figure 2E). The morphological defects of *mea* embryos are reminiscent of mutants with compromised embryonic patterning (Jenik et al., 2007; Möller and Weijers, 2009). To verify this hypothesis, we crossed *mea/MEA* with a set of tissue-specific markers for distinct embryonic domains (Figure 3A): apical-basal patterning (DR5V2 and *pPIN7::PIN7-GFP*, Figure 3B-D, Figure S3A,B), root tip architecture (*pPLT1::PLT1-YFP* and *pBBM::BBM-YFP*, Figure 3B,E-F, Figure S3C,D), provasculature (*pTMO5::3xGFP,* Figure 3B,G-H, Figure S3E), endodermis (*pSCR::SCR-GFP*, Figure 3B,i-J, Figure S3F), QC establishment (*pWOX5::dsRED,* Figure 3B,K-L, Figure S3G), and shoot apical meristem specification (SAM; *pWUS::dsRED* and *pCLV3::GFP,* Figure 3B,M-P, Figure S3H-I). We observed severely altered expression patterns of markers for different root regions, including expansion of the expression domain (*pPLT1::PLT1-YFP*, *pBBM:*:*BBM-YFP*, *pPIN7-PIN7::GFP*, Figure 3F, Figure S3C-D), ectopic expression in a different embryonic region (*DR5V2*, *pTMO5::3xGFP*, Figure 3D,H, Figure S3A,E), and absence of the signal suggesting loss of cellular identity (*pSCR::SCR-GFP* and *pWOX5::dsRED*, Figure 3J,L, Figure S3G). Consistent with the fact that cotyledons in *mea* embryos are only mildly affected, the shoot apical meristem domain was properly specified with only a small percentage of embryos showing weak or ectopic marker expression (*pWUS::dsRED* and *pCLV3::GFP,* Figure M-P, Figure S3H-I).

**Figure 2.**
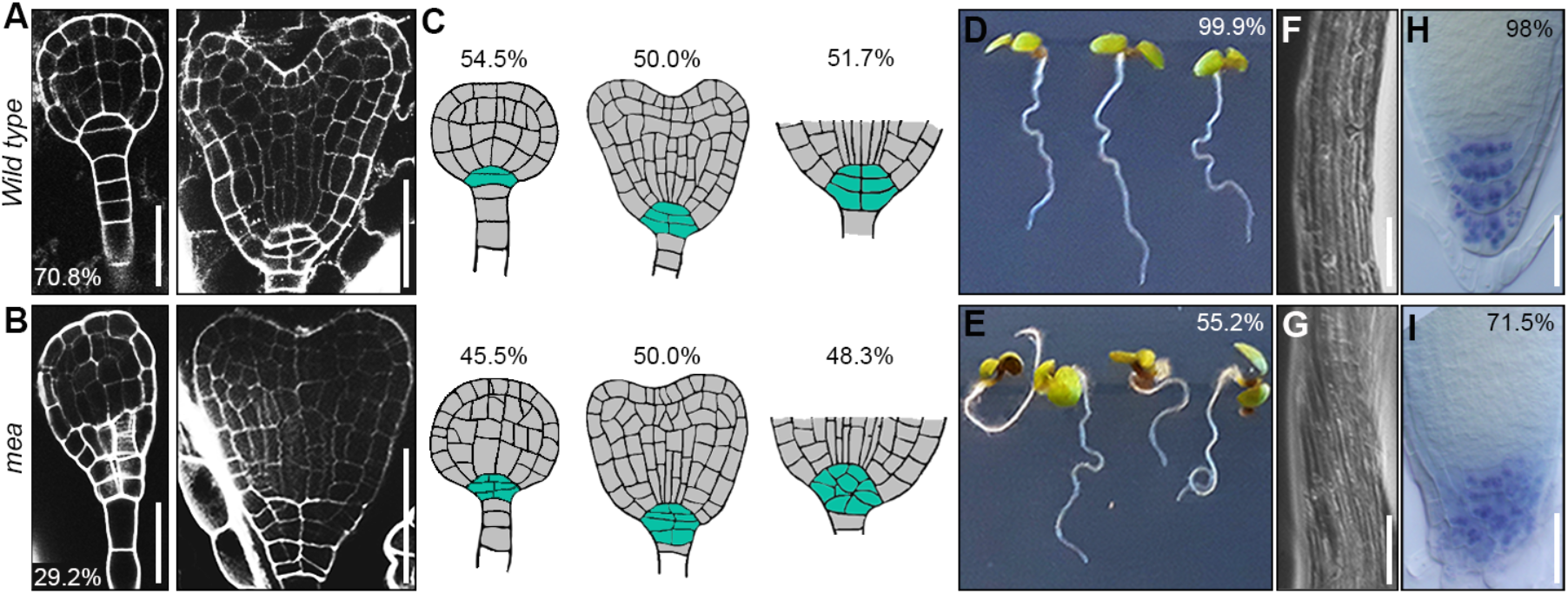
*mea* embryos develop severe morphological defects. **A-B.** mPS-PI staining of *mea/MEA* seeds showing wild-type (A) and aberrant (B) morphology at the globular (left image) and late heart stage (right image). **C.** Schematic representation of embryos in *mea/MEA* plants with wild-type (top row) or *mea* (bottom row) phenotypes obtained using mPS-PI images as template. Late globular, heart, and the basal part of late heart stage embryos are shown from left to right. The columella/QC region is highlighted in turquoise. **D-E.** Wild-type (E) and *mea* homozygous (F) seedlings grown on vertical plates. **F-G.** Magnification of epidermal cells of the primary root of wild-type (G) and *mea* homozygous (H) seedlings grown vertically. **H-I.** Lugol staining of the primary root tip of wild-type (I) and *mea* homozygous (J) seedlings. Scale bars: 25μm (A-B, left panels; H-J), 50μm (A-B, right panels), 250μm (F-G).

**Figure 3.**
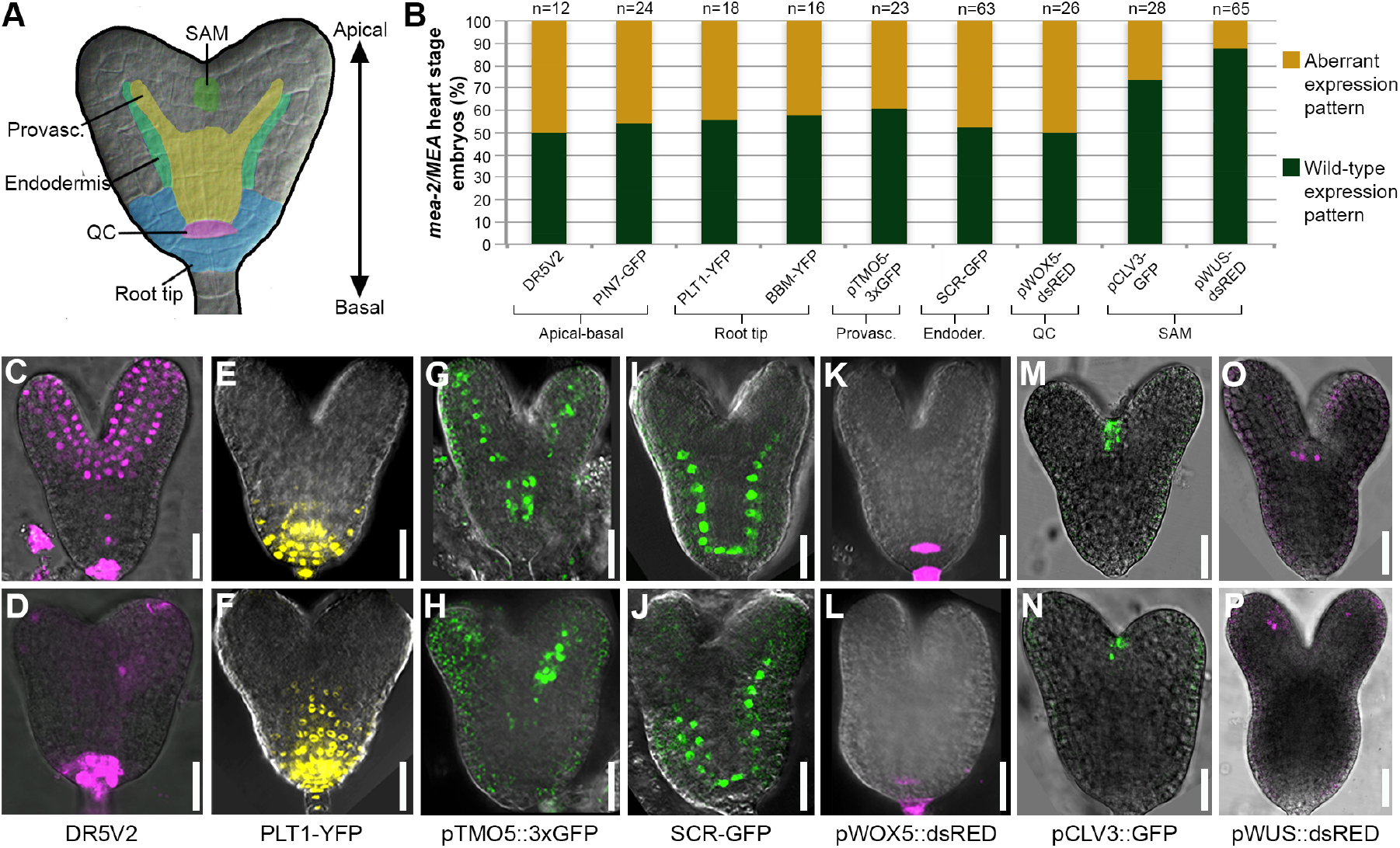
Embryonic patterning is affected in *mea* embryos. **A.** Schematic representation of the analyzed embryonic domains. **B.** Percentage of embryos showing altered expression pattern of the corresponding marker line in *mea/MEA* seeds. **C-P.** Confocal images showing the expression patterns in wild-type embryos (upper row) and *mea*-like (lower row) embryos at late heart stage for *DR5V2* (C-D), *pPLT1::PLT1-YFP* (E-F), *pTMO5::3xGFP* (G-H), *pSCR::SCR-GFP* (I-J), *pWOX5-dsRED* (K-L), *pCLV3::GFP* (M-N), and *pWUS::dsRED* (O-P). Scale bar: 20μm

In summary, from early stages onwards *mea* embryos display altered polarity, unbalanced symmetry, and an abnormal specification of embryonic domains and tissues. These results strongly suggest that *MEA* is required for proper embryonic patterning. Thus, in *Arabidopsis*, the spatial and temporal definition of the embryonic body plan relies on PcG proteins as it does in animals (Margueron and Reinberg, 2011).

### Cell cycle progression is compromised in *mea* embryos

We adopted a transcriptomic approach to identify the causative genes responsible for the patterning defects in *mea* embryos. Total RNA of *mea* homozygous and wild-type ovaries or seeds was collected at three time points: before fertilization (Ovary), one-two days after pollination (1-2DAP), and four days after pollination (4DAP). As absence of PRC2 activity leads to transcriptional de-repression (Kirmizis et al., 2004), we focused on genes showing significant up-regulation in *mea* mutants compared to the wild type (Supplementary Table 1). To enrich for potential targets responsible for the defects in *mea* embryos, we selected genes that were uniquely up-regulated in the 4DAP dataset (Figure 4A), the time-point when embryo-derived transcripts are technically detectable. After 138 up-regulated genes were represented in the 4DAP dataset only (Figure 4A), with 95 of them showing no detectable expression in wild-type embryos at the 16-cell (16C), early globular (EG), and late globular (LG) stages (INTACT datasets (Palovaara et al., 2017); Figure 4A and Supplementary Table 1). Manual annotation of molecular functions of these genes (represented as word-cloud, Figure 4A), revealed a predominant representation of factors involved in transcription, transport, enzymatic activity, hormone homeostasis, and the regulation of cell cycle (Figure 4A). We focused on the latter since *mea* embryos have more cells compared to the wild type (Grossniklaus et al., 1998). In order to measure the rate at which cells divide in *mea* embryos, we crossed *mea/MEA* plants with a triple cell cycle marker line (Desvoyes et al., 2020), enabling the simultaneous visualization of G1, S+early G2, and late G2+M phases. Two distinct classes of embryonic expression patterns were identified (Figure 4B-F, Figure S4): in the first, many nuclei were in G1 (high G1/G2 ratio, i.e. CFP-positive/RFP-positive nuclei), whereas the second had a low G1/G2 ratio. At heart and late heart stages, when *mea* embryos are clearly distinguishable from the wild type, all embryos with a low G1/G2 ratio displayed the *mea* phenotype (Figure.4E-F). As CFP is fused to the CTD1a protein, which is rapidly degraded upon entry into S-phase, a low number of CFP-positive nuclei indicates that more cells have entered S-phase and are thus committed to divide.

**Figure 4.**
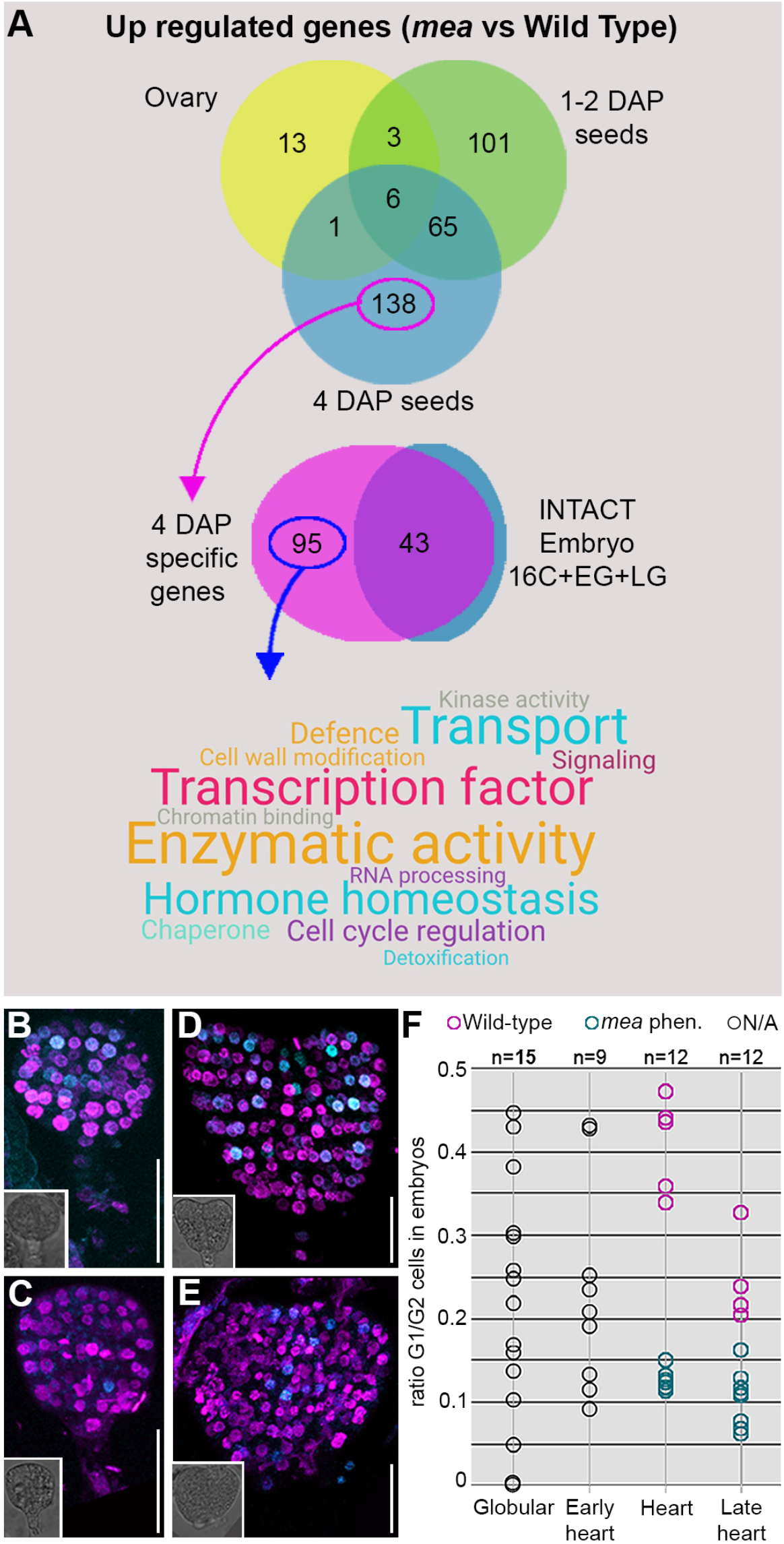
*mea* embryos display an accelerated cell cycle. **A.** Schematic representation of transcriptome analyses of *mea* homozygous versus wild-type ovaries/developing seeds. **B-E.** Confocal microscopy images of embryos of *mea/MEA* seeds expressing the triple cell-cycle marker line. Images show *CTD1-CFP* (G1) and *H3.1-RFP* (S+early G2) signals; M-phase marker is not included. Inlets in bottom left corners: brightfield images of the embryos analyzed. **F.** Quantification of the G1/G2 ratio in embryos of *mea/MEA* seeds. Pink circles are wild-type-looking, turquoise circles *mea*-like embryos, respectively; grey circles are embryos for which a phenotypic distinction was not possible. Scale bar: 20μm

Thus, the cell cycle progression through G1 is accelerated in *mea* embryos, a function that has also been described for maternally expressed imprinted genes in mammals (Barlow and Bartolomei, 2014; Lau et al., 1994; Leighton et al., 1995; Wang et al., 2007). In summary, our results support a role for *MEA* in regulating the embryonic body plan through the control of cell division.

### Deregulation of a core cell cycle component leads to *mea* embryonic defects

Accelerated and disorganized, ectopic cell divisions arise from the deregulation of cell cycle components (Gutierrez, 2009). D-type cyclins (CYCD) are conserved core constituents of the cell cycle machinery that integrate cell division and tissue patterning by promoting the G1-S transition (Meijer and Murray, 2000). Altered CYCD levels are sufficient to induce cell division by shortening the G1 phase and to trigger formative division defects (Forzani et al., 2014; Sozzani et al., 2010), a phenotype we observed in *mea* embryos. *CYCD1;1* was specifically up-regulated at 4DAP in *mea* seeds (Supplementary Table 1) and its increased expression was confirmed by ddPCR on RNA isolated from embryos around the early globular stage (Figure S5A). Visualization of a *pCYCD1;1::NLS-3xVenus-3’UTR* marker line revealed ectopic expression of *CYCD1;1* in the basal part of almond-shaped *mea* embryos (16.4%, n=104; Figure 5A, Figure S5A-C). Notably, the signal was undetectable in the endosperm and sibling wild-type embryos (Figure 5A), identifying CYCD1;1 as potentially responsible for the *mea* embryonic patterning defects. In line with this hypothesis, seed abortion was reduced from 50% in *mea/MEA* plants to 35.5% in *mea/MEA cycd1;1* double mutants (n=816; Figure 5B). Analysis of the F3 generation confirmed that the rescued seeds had inherited a maternal *mea* allele, with 8.0% of the progeny of *mea/MEA cycd1;1* plants being homozygous for *mea* (n=226; Figure 5B). Doubly homozygous *mea cycd1;1* plants exhibited 41.2% viable seeds (n=1883) of swollen and rounded appearance, *mea*-like endosperm, and mildly deformed giant embryos (Figure 5C-E; Figure S6A) with a significantly reduced number of cells in the columella/QC region as compared to *mea* (Figure 5F, Figure S6B-F). As a consequence of the less disorganized root meristem, *mea cycd1-1* seedlings showed restoration of the primary root’s gravitropic response (Figure 5G, Figure S7A). These results unequivocally demonstrate that the removal of *CYCD1;1* activity is sufficient to rescue *mea* embryos and leads to a bypass of growth arrest even if they are surrounded by abnormal *mea* endosperm.

**Figure 5.**
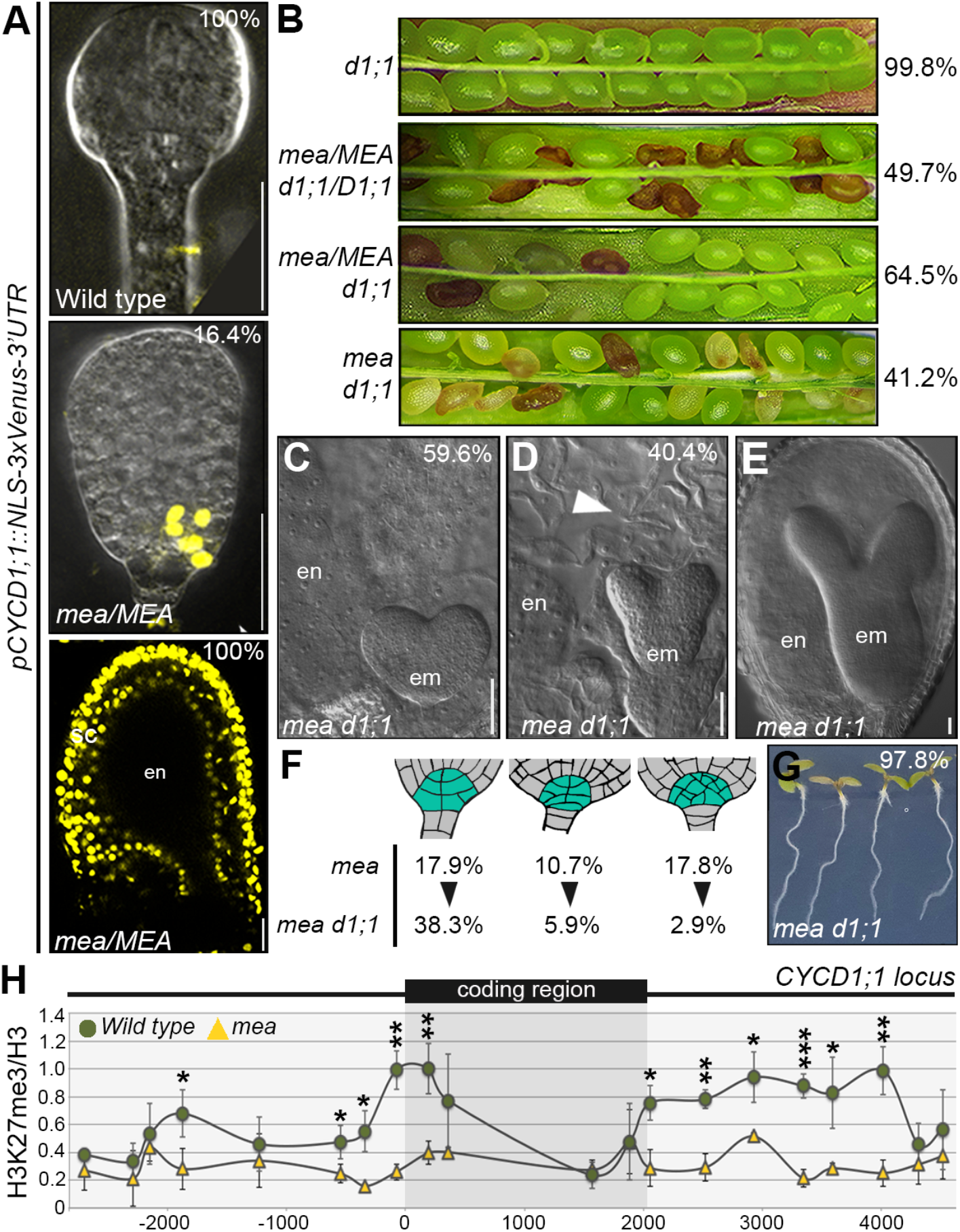
MEA patterns the embryo through regulation of a core cell cycle component. **A.** Confocal microscopy images of embryos of *mea/MEA* seeds showing expression of *pCYCD1;1::NLS-3xVenus-3’UTR* in wild-type (left) and *mea* (middle) embryos, and *mea*/*MEA* seeds. **B.** Opened siliques (from top to bottom): *cycd1;1*, *mea/MEA cycd1;1/CYCD1;1, mea/MEA cycd1;1*, and *mea cycd1;1* plants with percentage of viable seeds indicated on the right. **C-E.** DIC microscopy images of seeds from *mea cycd1;1* plants showing a *mea*-like seed (C), a seed with a wild-type-looking embryo surrounded by *mea*-looking endosperm (D), and a seed with a giant embryo (E). Arrowhead indicates uncellularized endosperm. **F.** Schematic representation of cell number and organization in the columella/QC region in *mea cycd1;1* embryos compared to *mea*. The number of embryos showing a given range of cells is represented as a percentage. **G.** Gravitropic response of vertically grown *mea cycd1;1* seedlings. **H.** CUT&RUN analysis of H3K27me3 over H3 occupancy at the *CYCD1;1* locus in wild-type versus *mea* embryos. Error bars: Standard deviation. P value of a *t*-test: * <0.05, **<0.01, ***<0.001. em, embryo; en, endosperm. Scale bar: 20μm (A), 50μm (C-E)

In agreement with *CYCD1;1* being able to impose abnormal embryonic cell divisions, ectopic expression of *CYCD1;1* in embryos induced seed abortion ranging from 5 to 20% (*pRPL18::CYCD1;1;* Figure S7B,C), with embryos arrested at the late globular stage and exhibited proliferation defects in the basal part (13.8%, n=894, Figure S7C), reminiscent of almond-shaped *mea* embryos at a similar stage (Figure 2B). Furthermore, 31.2% of the viable embryos showed aberrant division planes in the root meristem (n=461, Figure S7D). Remarkably, albeit *pRPL18* driving expression also in the endosperm (Yan et al., 2016) (Figure S7E), none of the 18 *pRPL18::CYCD1;1* lines analyzed showed abnormal endosperm proliferation and/or cellularization (Figure S7C). These results show that ectopic expression of *CYCD1;1* is sufficient to phenocopy the patterning defects observed in *mea* embryos, independent of the genotype of the endosperm.

To determine whether *CYCD1;1* is indeed a direct FIS-PRC2 target, we analyzed H3K27me3 levels at the *CYCD1;1* locus (coding region plus 2.5 kb up- and downstream sequences) in isolated early globular *mea* and wild-type embryos by CUT&RUN (Figure 5H). Consistent with the increased expression of *CYCD1;1*, H3K27me3 at the *CYCD1;1* locus was significantly reduced in *mea* embryos compared to the wild type, in particular around the transcriptional start site and downstream of the coding region but also in a region about 1.8kb upstream.

Taken together, our results show that *CYCD1;1* is a key target of maternal *MEA* and that its deregulation in *mea* embryos is majorly responsible for their abnormal development and abortion. Thus, PRC2 directly regulates embryonic patterning and growth via the repression of *CYCD1;1*, a core cell cycle component.

## DISCUSSION

PcG proteins are central to both animal and plant development (Inoue et al., 2017; Raissig et al., 2013), but through which target genes they exert this control is known for only a few plant developmental processes (Goodrich et al., 1997; Ikeuchi et al., 2015; Köhler et al., 2003b; Lodha et al., 2013). Although mutations affecting FIS-PRC2 cause maternal effect embryo abortion (Grossniklaus et al., 1998), a direct role of PcG proteins in plant embryogenesis has been dismissed(Bouyer et al., 2011; Kiyosue et al., 1999; Leroy et al., 2007; Scott et al., 1998). We show that, independent of the genotype of the endosperm, *mea* embryos develop severe patterning defects as a result of abnormal cell divisions, clearly demonstrating a direct role of FIS-PRC2 in embryonic patterning. Overproliferation in *mea* embryos is caused by derepression of the core cell cycle component *CYCD1;1*, known to promote the rate and direction of cell divisions (Forzani et al., 2014; Meijer and Murray, 2000) but is usually silenced in the early embryo by H3K27me3 mediated by MEA. *CYCD1;1* is a major target of MEA as its overexpression in the wild type causes defects reminiscent of those observed in *mea* embryos and the *cycd1;1* mutant largely suppresses the *mea* phenotypes. As suppression of *mea* seed abortion is incomplete, additional MEA target genes may play a minor role in embryonic patterning.

The PRC2 regulatory complex is conserved from animals to plants^1^ and thus arose in their common, unicellular ancestor before the split of the two kingdoms. Interestingly, PRC2 regulates cell proliferation and pattern formation not only in plants as shown here but also in animals (O’Carroll et al., 2001; O’Dor et al., 2006; Oktaba et al., 2008; Pasini et al., 2004), despite the fact that multicellularity evolved independently in these lineages. Similarly, genomic imprinting arose through convergent evolution in plants and mammals but does exert growth control and is partly regulated by PRC2 in both these groups (Barlow and Bartolomei, 2014; Ferguson-Smith, 2011; Grossniklaus and Paro, 2014; Pires and Grossniklaus, 2014). It is possible that PRC2 had an ancient role in regulating the cell cycle in the common ancestor of animals and plants and that this module was then exploited as pattern formation evolved in multicellular organisms and again as a placental habit and genomic imprinting arose in seed plants and mammals, respectively (Inoue et al., 2017; Pires and Grossniklaus, 2014). Thus, the regulation of cell proliferation by PRC2 seems to form a robust module that was independently recruited into various epigenetically controlled processes during the evolution of multicellular organisms.

## Acknowledgements

We thank Dolf Weijers and Ben Scheres (Wageningen University and Research) and the Nottingham Arabidopsis Stock Center for providing seeds, Richard Immink (Wageningen University and Research) and Dorus Gadella (University of Amsterdam) for providing elements of the TagRFP destination vector, Thomas Laux (Albert-Ludwigs-Universität Freiburg), in whose laboratory pWOX5::DsRED was generated, Steve Henikoff (Fred Hutchinson Cancer Research Center) for the gift of pA-Mnase, spike-in DNA, and helpful suggestions on CUT&RUN, Christof Eichenberger, Frédérique Pasquer, Arturo Bolaños, Daniela Guthörl, and Peter Kopf (University of Zurich) for general lab support, and Gregor Rot (University of Zurich) for technical assistance during the RNA-Seq raw data deposition. We thank Andrea Steimer (University of Zurich) for sharing the pRPS5a::MEA construct prior publication. Funding: This work was supported by the University of Zurich, postdoctoral fellowships from Europen Molecular Biology Organization and the European Union’s Marie Skłodowska-Curie Action to MB and SB, respectively, grant RTI2018-094793-B-I00 from Ministerio de Ciencia e Innovación (MICIN-FEDER) to CG, and grants from the Swiss National Science Foundation, the 5^th^ Framework Program of the European Union, and the European Research Council to UG.

## Author contributions

UG conceived and supervised the project. SS designed and performed most of the experiments. SS and UG analyzed and interpreted the data. MB created the double pollination lines and performed RNA-Seq together with NP. SB generated CYCD1;1 marker and overexpression lines, AS made the MEArescue line, and BD and CG generated and provided the triple cell cycle marker line. VG performed ddPCR on material collected by SS. EvdG generated the pWOX5::DsRed marker line. SS and UG wrote and all authors commented on the manuscript.

## Authors’ approval

All authors have read and approved the manuscript.

## Competing interests

All authors declare that there are no competing interests.

## METHODS

### Plant material and growth conditions

Seeds were sown on half-strength MS media (1/2 MS salt base [Carolina Biologicals, USA], 1% sucrose, 0.05%MES, 0.8% Phytoagar [Duchefa], pH>5.7 with KOH), stratified for 3-4 days at 4°C in the dark, and then moved to long-day conditions (8h dark at 18°C, 16h light at 22°C, 70% humidity). When showing four true leaves, seedlings were transplanted to soil and grown under long-day conditions in a walk-in growth chamber (8h dark, 16h light, 22°C, 70% humidity). Lines used in this study are: *DR5V2*(Liao et al., 2015), *pPIN7::PIN7-GFP*(Vieten et al., 2005) (NASC ID N9577), *pPLT1::PLT1-YFP*(Galinha et al., 2007), *pBBM::BBM-YFP*(Galinha et al., 2007), *pTMO5::3xGFP*(Schlereth et al., 2010), *pSCR::SCR-GFP*(Wysocka-Diller et al., 2000) (NASC ID N3999), *pCLV3::GFP-ER pWUS::dsRED-N7*(Gordon et al., 2007) (NASC ID N23895); *cycd1-1* mutant (GABI_214D10); *pMEA::MEA-GR*(Pires et al., 2016); PlaCCI triple cell-cycle marker line(Simonini et al., 2016).

*mea* mutants used in this study: *mea-1* and *mea-2*(Grossniklaus et al., 1998) marked by the Kanamycin resistance gene. When required, progeny of crosses with *mea* were sown on Kanamycin half-strength MS plates to select for *mea/MEA* individuals. The various marker lines were introduced into the *mea/MEA* mutant background by crossing, using *mea/MEA* as pollen donor.

### Creation of *kpl-GFP*, *MEArescue-RFP* lines and double pollination

The *pRPS5A-GFP* construct was assembled as follow: the *pRPS5a* promoter was cloned into pDONR207 using Gateway cloning and subsequently into the pMDC107 destination vector(Curtis and Grossniklaus, 2003).

The *pRPS5A-TagRFP* was generated by Gateway cloning of the *pRPS5a* promoter into pDONR221 and subsequently in destination vector CZN654 (based on pB7WG2(Karimi et al., 2002) but adapted by Richard Immink with TagRFP, a gift from Dorus Gadella).

Flowers around stage 12 of spontaneous *mea-2* homozygous individuals or not DEX-induced *mea-1 pMEA::MEA-GR*(Pires et al., 2016) individuals were emasculated and pollinated 24h later. The first pollination was a minimal pollination with *kpl*-GFP pollen. The second pollination was done with *MEA-rescue+RFP* pollen 4.5h after the first pollination. This timing was chosen based on the speed at which pollen tubes grow in the pistils under our growth conditions, and evaluated by the rate of synergid rupture in ovules mounted in 7% glucose supplemented with 0.1mg/ml of Propidium Iodide (SIGMA, P4170) imaged by a Leica SP5microscope (Argon laser, excitation 488nm). Under our growth conditions, about 50% of ovules displayed synergid rupture 3.5h after pollination. Each set of single and double pollination experiment was performed three to five times, with similar results.

### Creation of *pWOX5::dsRED* line

The WOX5 promoter fragment(Sarkar et al., 2007) [1] was cloned from the AKS32 plasmid as Pst1 fragment into the ML939 cloning vector digested with Pst1, resulting in pEG126. The dsREDer reporter with 35s CaMV terminator sequence was cloned from the ML878 plasmid after SphI digest, blunting with T4 DNA polymerase and second digest with XhoI, into pEG126 digested with SmaI and SalI, resulting in pEG279. The pWOX5:dsREDer expression cassette was cloned from pEG279 with Pac1 and Asc1 into the binary plasmid ML516 digested with PacI and AscI, resulting in pEG280.

The vectors were introduced into *Agrobacterium tumefaciens* strain GV3101, and Wild type Col-0 plants transformed following the floral dip method(Clough and Bent, 1998). At least 20 independent transgenic lines were generated and examined for expression pattern, using the pWOX5:GFP line as control. The line analyzed here is T4 generation homozygous lines with medium/high expression. Ectopic expression observed outside the QC area is a direct consequence of the ability of the dsRED protein to make multimers that can be highly stable. Since the QC cells can divide to replace dead stem cells bordering the QC, sometime dsRED signal can be seen in areas neighboring the QC, in our analyses the upper cells of the suspensor. This ectopic signal observed in suspensor cells was excluded from the comparison between wild-type and aberrant embryos.

### Clearing

Siliques were fixed o/n in fixative (Ethanol:Acetic Acid 9:1 v/v) at room temperature. The following day, the fixative was replaced with Ethanol70%. Seeds were isolated from the valves and mounted in Hoyer’s solution (Chloral Hydrate:Water:Glycerol 10:2,5:1 w/v/w) and left to clear overnight. Small seeds required only a few hours of clearing. Images were taken with a Leica DM6000B or Zeiss DMR microscope, both equipped with differential interference contrast (DIC) filters and ANDOR 5.5 Neo sCMOS cameras.

### Lugol staining

Seedlings seven days after germination and grown on vertical plates were incubated for 2min in Lugol solution (Sigma 62650), rinsed in water, mounted in clearing solution (Chloral Hydrate:Water:Glycerol 8:4:1 w/v/w), and imaged immediately. Images were taken with a Zeiss DMR microscope equipped with DIC filters and an ANDOR 5.5 Neo sCMOS camera.

### Cloning of *CYCD1;1* marker and overexpression lines

The *pCYCD1;1-NLS-3xVenus-3’UTR* marker line includes the promoter region (5476 bp upstream of the ATG) and the 3’UTR (4349 bp downstream of the stop codon) of the *CYCD1;1* locus (*At1g70210*). The promoter and 3’UTR fragment were assembled as Golden Gate module together with the NLS-localization signal and the 3X-Venus, in a pPZP222 vector modified to accept Golden Gate modules(Bencivenga et al., 2016).

For the *pRPL18::CYCD1;1* transgenic line, the RPL18(Yan et al., 2016) promoter was cloned as Golden Gate module upstream of the *CYCD1;1* coding sequence (CDS, no introns), in the Golden Gate acceptor version of pPZP222; as terminator, the *35S*terminator was placed downstream the *CYCD1;1* CDS.

For the *pRPL18::NLS-3xVenus*, the module for pRPL18, NLS, 3xVenus, *35S*terminator were assembled in the pPZP222 vector.

Where necessary, site-specific mutagenesis was used to remove endogenous BsaI sites.

All vectors were introduced into *Agrobacterium tumefaciens* strain GV3101, and plants transformed following the floral dip method(Clough and Bent, 1998). The *pCYCD1;1-NLS-3xVenus-3’UTR* transgene was then introduced from a selected line into the *mea-2/MEA* background by crossing.

Primers are listed in the Extended Table 2.

### Droplet Digital PCR (ddPCR)

Genotyping of single seeds: individual seeds were removed from the fruit, deposited in a 1.5ml Eppendorf tube, and grinded with a blue plastic pestle. DNA was extracted with the Mag-Bind Plant DS DNA kit (OMEGA Bio-Tek) following the manufacturer’s instructions. Elution was done with water into a 1.5ml low-binding Eppendorf tube. The DNA samples were then concentrated in a Speedvac to 20μl final volume. Total DNA was digested for 30min at 37°C with EcoRV, and then pre-amplified with specific primers for GFP (GFP-fw + GFP-rev), RFP (RFP-fw + RFP-rev), and internal control (Control-fw + Control-rev) genes altogether in the same reaction tube, using the ExTaq linear polymerase (Takara, RR001A) as follows: DNA 3μl, 40nM each primer, 0.6Unit Extaq in 1x buffer containing 1.5mM MgCl_2_ with the following PCR protocol: 98°C × 3min, [98°C × 10sec; 59°C × 20sec; 72°C × 20sec]×15 cycles. For ddPCR, 5μl of pre-amplified DNA were used in duplex assays GFP/Control and RFP/Control. Assay conditions: 500nM primers, 200nM probes, and QX200 ddPCR Probe no dUTPS Supermix (BioRad). The PCR protocol was the manufacturer’s recommendation for Probe assays (95°C: 10min, 94°C 30s, Ramp 2.5°C/s, 60°C 1min, Ramp 2.5°C/s[40 cycles], 98°C 10min, 4°C until further process). Fluorescence was detected with the QX200 droplet digital reader (Bio-Rad), and analyzed with the provided Quanta Soft version 1.7 software. Presence of each gene was calculated relative to the endogenous control. The sum was given as 100% and ratio of each gene was calculated as % relative to total.

Expression analyses of *CYCD1;1* on isolated embryos: seeds were removed from siliques, placed in a 1.5ml Eppendorf tube containing PBS1X, and gently pressed with a blue plastic pestle with up-and-down movements to release the embryos. Collection time did not exceed 15min. The sample was then passed through a 100μm pore-size cell strainer (CellTrics) to remove excess of debris. The flow-through, containing the embryos, was collected in a small plastic rectangular box with low walls (we used the lid of the 8-well Tissue Culture Chambers REF94.6190.802, SARSTEDT). Embryos were collected with a 50μm diameter size capillary (ES-blastocyst injection pipettes, BioMedical Instruments, BM100T-10P) and an oil micromanipulator (CellTram Vario, Eppendorf), mounted on a Leica SP2 inverted microscope. Embryos were collected in maximum 1h shifts and washed thoroughly in fresh PBS1X. The drop of PBS1X buffer containing the embryos was then ejected directly from the capillary onto a piece of parafilm to create a round-shaped droplet. The parafilm was then placed for 2min at −70°C to let the droplet freeze. Frozen droplets were collected in a 1.5ml low-binding Eppendorf tube and stored at −70°C until the extraction. For our experiment, a total of 1000 embryos around the early globular stage (with and without suspensor) were collected per replicate from wild-type and *mea* plants, and ddPCR was performed on biological triplicates (total of 3000 embryos per genotype). For RNA extraction, 4-6 glass beads (1.7-2.1mm diameter, ROTH A556-1) were added to each tube containing the frozen droplets with the embryos, frozen in liquid nitrogen, and grinded 3-4 times with a single-tube tissue grinder (Silamat S6). The RNA extraction was done with the Qiagen RNeasy Plant Mini extraction kit, and subsequently treated with Turbo-Dnase (Ambion) following the manufacturer’s protocol. cDNA synthesis was performed using the Maxima Reverse Transcriptase (Invitrogen) and OligodT (Invitrogen) following the manufacturer’s protocol. 5μl of a 1:2 dilution of cDNA were then used for ddPCR assays *CYCD1;1/UBI21*, with 100nM final concentration of each primer, in a total reaction volume of 25μl, 20 of which were used to generate droplets in 1X Master mix EVAGREEN (BIORAD). PCR conditions were according to manufacturer’s recommendation for EVAGREEN.

Primers are listed in the Extended Table 2.

### RNA-SEQ

The following tissue was harvested for the three different stages: i) ovaries two days after emasculation (style and stigma were removed), ii) developing seeds 1-2 days after pollination (dissected from siliques), and iii) developing seeds 4 days after pollination (dissected from siliques). Three independent biological replicates were generated for each tissue/genotype combination. For each replicate, the isolated tissue were frozen in liquid nitrogen, ground to a fine powder using a pestle and incubated for 10 minutes in 450 μl of a solution containing 2% CTAB, 100 mM Tris-HCl pH 8.0, 25 mM EDTA, 2M NaCl and 2% ß-mercaptoethanol. This suspension was then mixed with ice-cold chloroform was centrifuged 15 minutes at 16000 g. The upper phase was collected and mixed with 150μl of a 8M LiCl solution, incubated at −20 °C for one hour and centrifuged 30 minutes at 16000 g. The RNA pellet was washed with 70% ethanol and resuspended in 30 μl RNAse free water and quantified using the Qubit. The Turbo DNA free kit (Ambion AM1907) was used to remove DNA. Total RNA samples were polyA enriched and reverse-transcribed into double stranded cDNA. Sequencing libraries were generated using the TruSeq RNA Sample Prep Kit v2 (Illumina). Libraries were normalised and pooled using TruSeq index adapters and sequenced using single reads in a Illumina HiSeq 2000 sequencer at the Functional Genomics Centre Zurich. Low quality read ends were clipped and reads were mapped to the TAIR reference genome with TopHat. Differential gene expression was performed using DEseq2. For the differential gene expression analysis between wild-type and *mea* samples, only genes for which more than 4 counts per million were present in most samples were analysed. After dispersion estimates were obtained, a negative binomial model was fitted and differential expression was tested using a quasi-likelihood F-test, as implemented in EdgeR (using a p-value of 0.01)

GO analyses was performed with g:Profiler (http://biit.cs.ut.ee/gprofiler/gost).

### Confocal imaging, mPS-PI, and PI staining

Confocal analyses are done using a Leica SP5 confocal microscope. GFP: Argon laser, excitation 488; YFP: Argon laser, excitation 514nm; CFP: argon laser, excitation 456nm; RFP: argon laser excitation 558nm; Propidium iodide: Argon laser, excitation 488.

mPS-PI of seeds: seeds at different developmental stages were isolated from siliques and treated as in Truernit et al., 2008.(Truernit et al., 2008)

PI staining of primary root: seedlings were incubated for 10min in Propidium Iodide solution at a concentration of 10μg/ml (Sigma P4170), mounted in glycerol 30%, and imaged with a Leica SP5, Argon Laser, excitation 488nm.

### CUT&RUN

Embryos for CUT&RUN were isolated as described above for the expression analysis of *CYCD1;1*, with the exception that the collection buffer PBS1X was supplemented with the Mini Protease EDTA-free Inhibitor Cocktail (ROCHE). A total of 3500 embryos were collected around the globular stage per replicate per genotype (wild-type and *mea*). CUT&RUN has been performed in triplicate following the protocols reported by Zheng and Gerhing, 2019 and Skene et al., 2018, with some minor modifications. Differently from Zheng and Gerhing, 2019, we included Tween-20 in the binding and washing buffers as in Skene et al., 2018. Antibodies used were: anti-H3K27me3 (Abcam 192985) and anti-H3 (Abcam 1971). pA-Mnase and the Spike-in DNA were a kind gift of Steve Henikoff. The final pellet of DNA was resuspended in 50μl of TE1X. Real-Time PCR was performed on a BIORAD CFX384 machine, in a technical triplicate, on 384-well plates (LightCycler 480 white plates and sealing foils, ROCHE), using 0.7μl of DNA per replicate and the SsoAdvanced Universal SYBR Green Supermix (BIORAD). Results are presented as H3K27me3 enrichment over H3 occupancy. Statistical analyses are based on a *t*-test. Primers are listed in the Extended Table 2.

## SUPPLEMENTARY INFORMATION

**Supplementary Figure 1.**

**A.** Quantification of phenotypes observed in single and double crosses.

**B.** Images of cleared seeds derived from the double pollination of *mea* pistils. Wild-type embryo is surrounded by uncellularized endosperm.

**C.** Appearance of developing seeds (up) and dried seeds (bottom) in double pollination of *mea* pistils. The developing embryo (arrow head) is visible in big and swollen seeds, a phenotype that is characteristic of *mea* seeds. Small seeds are the ones derived from full rescue by the *MEA-rescue+RFP* pollen (asterisk).

**D.** ddPCR analyses for presence of *GFP/RFP* transgenes in single seeds derived from the double pollination of *mea* pistils.

**Supplementary Data Figure 2.**

**A.** Quantification of the number of embryos showing aberrant division planes in *mea/MEA* heterozygous offspring.

**B.** mPS-PI staining of *mea/MEA* seeds, showing examples of embryos from the globular (left) to the late heart (right) stages.

**C.** Quantification of the number of cells in the columella/QC region in embryos at late globular stage showing a wild-type phenotype or aberrant divisions.

**D.** Quantification of the number of cells in the columella/QC region in embryos at late heart stage showing a wild-type or *mea* phenotype.

**E.** Lugol staining and Propidium Iodide staining of root tips of wild-type and *mea* homozygous seedlings. The percentage shows the frequency of the phenotype observed in the imaged seedlings (n=14).

**Supplementary Data Figure 3.**

For all markers, histograms of the quantification are shown on the left and pictures of representative embryos on the right; pictures show wild-type expression patterns in the top row and aberrant ones in the bottom row for all markers, except for (b) where wild-type and aberrant expression patters are shown in the left and right panels, respectively.

**A.** Quantification of aberrant versus wild-type expression patterns in *mea/MEA DR5V2* seeds.

**B.** Quantification of aberrant versus wild-type expression patterns in *mea/MEA pPIN7::PIN7-GFP* seeds.

**C.** Quantification of aberrant versus wild-type expression patterns in *mea/MEA pPLT1::PLT1-YFP* seeds.

**D.** Quantification of aberrant versus wild-type expression patterns in *mea MEA pBBM::BBM-YFP* seeds.

**E.** Quantification of aberrant versus wild-type expression pattern in *mea/MEA pTMO5::3xGFP* seeds.

**F.** Quantification of aberrant versus wild-type expression patterns in *mea/MEA pSCR::SCR-GFP* seeds.

**G.** Quantification of aberrant versus wild-type expression patterns in *mea/MEA pWOX5::dsRED* seeds.

**H.** Quantification of aberrant versus wild-type expression patterns in *mea/MEA pCLV3::GFP* seeds.

**I.** Quantification of aberrant versus wild-type expression patterns in *mea/MEA pWUS::dsRED* seeds.

**Supplementary Data Figure 4.**

**A.** Confocal images of embryos in *mea/MEA* seeds. Turquoise signal is CTD1-CFP marking G1; magenta signal is H3.1-RFP marking S+early G2. M-phase marker is not included.

**Supplementary Data Figure 5.**

**A.** ddPCR analyses of *CYCD1;1* transcript level in wild-type versus *mea* embryos around early globular stage.

**B.** Percentage of embryos from *mea/MEA* seeds showing altered *pCYCD1;1::NLS-3xVenus-3’UTR* expression patterns.

**C.** Confocal images of *pCYCD1;1::NLS-3xVenus-3’UTR* marker line showing its wild-type expression pattern in embryos from the globular to mature stages, a magnification of the root tip showing no expression in the QC, and a seed showing expression only in the sporophytic tissues.

**D.** Confocal images of *pCYCD1;1::NLS-3xVenus-3’UTR* marker line showing its aberrant expression in *mea/MEA* seeds, in embryos from the globular to late heart stages. In *mea/MEA* seeds, no expression of the marker is detected in the endosperm.

**Supplementary Data Figure 6.**

**A.** DIC analyses of *mea cycd1;1* seeds, showing enlarged embryos surrounded by defective endosperm.

**B-E.** Quantification of the number of cells in the columella/QC region in embryos at the globular (b), early heart (c), heart (d), and late heart (e) stages of wild-type and *cycd1;1*, *mea/MEA*, *cycd1;1/1CYCD1;1 mea/MEA* doubley heterozygous, and *cycd1;1 mea/MEA* mutants.

**F.** mPS-PI staining of *mea/MEA cycd1;1* embryos.

**Supplementary Data Figure 7.**

**A.** Lugol staining of root tip of *mea cycd1;1* homozygous seedlings.

**B.** Confocal analyses of *pRPL18::NLS-3xVenus* showing expression in embryos and endosperm.

**C.** Images of rosettes *of pRPL::NLS-3xVenus* (left) and embryo at globular stage by DIC analyses (right).

**D.** Images of rosettes *of pRPL18::CYCD1;1* (left), seed abortion (middle) and by DIC analyses of seeds (right), showing embryos with abnormal shape surrounded by normal endosperm.

**E.** mPS-PI staining of *pRPL18-CYCD1;1* seeds showing embryos with proliferation in the basal part.

**Supplementary Table 1.**

Contains:

- Up-regulated and Down-regulated genes in *mea* vs Wild-type RNA-Seq of Ovary tissue.
- Up-regulated and Down-regulated genes in *mea* vs Wild-type RNA-Seq of 1-2DAP seeds.
- Up-regulated and Down-regulated genes in *mea* vs Wild-type RNA-Seq of 4DAP seeds.
- Overlap 4DAP specific dataset with INTACT dataset of 16C, EG and LG globular embryos.

**Supplementary Table 2.**

Primers used in this study.

## Notes

### Competing Interest Statement

The authors have declared no competing interest.

## REFERENCES

Barlow, D.P., Bartolomei, M.S., 2014. Genomic Imprinting in Mammals. Cold Spring Harb. Perspect. Biol. 6, a018382. https://doi.org/10.1101/cshperspect.a018382

Bencivenga, S., Serrano-Mislata, A, Bush, M., Fox, S., Sablowski, R., 2016. Control of Oriented Tissue Growth through Repression of Organ Boundary Genes Promotes Stem Morphogenesis. Dev. Cell 39, 198–208. https://doi.org/10.1016/j.devcel.2016.08.013

Bouyer, D., Roudier, F., Heese, M., Andersen, E.D., Gey, D., Nowack, M.K., Goodrich, J., Renou, J.-P., Grini, P.E., Colot, V., Schnittger, A., 2011. Polycomb Repressive Complex 2 Controls the Embryo-to-Seedling Phase Transition. PLOS Genet. 7, e1002014. https://doi.org/10.1371/journal.pgen.1002014

Chanvivattana, Y., Bishopp, A., Schubert, D., Stock, C., Moon, Y.-H., Sung, Z.R., Goodrich, J., 2004. Interaction of Polycomb-group proteins controlling flowering in Arabidopsis. Development 131, 5263–5276. https://doi.org/10.1242/dev.01400

Chaudhury, A.M., Ming, L., Miller, C., Craig, S., Dennis, E.S., Peacock, W.J., 1997. Fertilization-independent seed development in Arabidopsis thaliana. Proc. Natl. Acad. Sci. U. S. A. 94, 4223–4228.

Clough, S.J., Bent, A.F., 1998. Floral dip: a simplified method forAgrobacterium-mediated transformation ofArabidopsis thaliana: Floral dip transformation of Arabidopsis. Plant J. 16, 735–743. https://doi.org/10.1046/j.1365-313x.1998.00343.x

Curtis, M.D., Grossniklaus, U., 2003. A Gateway Cloning Vector Set for High-Throughput Functional Analysis of Genes in Planta. Plant Physiol. 133, 462–469. https://doi.org/10.1104/pp.103.027979

Desvoyes, B., Arana-Echarri, A, Barea, M.D., Gutierrez, C., 2020. A comprehensive fluorescent sensor for spatiotemporal cell cycle analysis in Arabidopsis. Nat. Plants 1–5. https://doi.org/10.1038/s41477-020-00770-4

Faust, C., Schumacher, A., Holdener, B., Magnuson, T., 1995. The eed mutation disrupts anterior mesoderm production in mice. Development 121, 273–285.

Ferguson-Smith, A C., 2011. Genomic imprinting: the emergence of an epigenetic paradigm. Nat. Rev. Genet. 12, 565–575. https://doi.org/10.1038/nrg3032

Förderer, A., Zhou, Y., Turck, F., 2016. The age of multiplexity: recruitment and interactions of Polycomb complexes in plants. Curr. Opin. Plant Biol. 29, 169–178. https://doi.org/10.1016/j.pbi.2015.11.010

Forzani, C., Aichinger, E., Sornay, E., Willemsen, V., Laux, T., Dewitte, W., Murray, J.A.H., 2014. WOX5 suppresses CYCLIN D activity to establish quiescence at the center of the root stem cell niche. Curr. Biol. CB 24, 1939–1944. https://doi.org/10.1016/j.cub.2014.07.019

Galinha, C., Hofhuis, H., Luijten, M., Willemsen, V., Blilou, I., Heidstra, R., Scheres, B., 2007. PLETHORA proteins as dose-dependent master regulators of Arabidopsis root development. Nature 449, 1053–1057. https://doi.org/10.1038/nature06206

Goodrich, J., Puangsomlee, P., Martin, M., Long, D., Meyerowitz, E.M., Coupland, G., 1997. A Polycomb-group gene regulates homeotic gene expression in Arabidopsis. Nature 386, 44–51. https://doi.org/10.1038/386044a0

Gordon, S.P., Heisler, M.G., Reddy, G.V., Ohno, C., Das, P., Meyerowitz, E.M., 2007. Pattern formation during de novo assembly of the Arabidopsis shoot meristem. Development 134, 3539–3548. https://doi.org/10.1242/dev.010298

Grossniklaus, U., Paro, R., 2014. Transcriptional Silencing by Polycomb-Group Proteins. Cold Spring Harb. Perspect. Biol. 6, a019331. https://doi.org/10.1101/cshperspect.a019331

Grossniklaus, U., Vielle-Calzada, J-P., Hoeppner, M.A., Gagliano, W.B., 1998. Maternal Control of Embryogenesis by MEDEA, a Polycomb Group Gene in Arabidopsis. Science 280, 446–450. https://doi.org/10.1126/science.280.5362.446

Gutierrez, C., 2009. The Arabidopsis Cell Division Cycle. Arab. Book Am. Soc. Plant Biol. 7. https://doi.org/10.1199/tab.0120

Ikeuchi, M., Iwase, A., Rymen, B., Harashima, H., Shibata, M., Ohnuma, M., Breuer, C., Morao, A.K., de Lucas, M., De Veylder, L., Goodrich, J., Brady, S.M., Roudier, F., Sugimoto, K., 2015. PRC2 represses dedifferentiation of mature somatic cells in Arabidopsis. Nat. Plants 1, 15089. https://doi.org/10.1038/nplants.2015.89

Inoue, A., Jiang, L., Lu, F., Suzuki, T., Zhang, Y., 2017. Maternal H3K27me3 controls DNA methylation-independent imprinting. Nature 547, 419–424. https://doi.org/10.1038/nature23262

Jenik, P.D., Gillmor, C.S., Lukowitz, W., 2007. Embryonic Patterning in Arabidopsis thaliana. Annu. Rev. Cell Dev. Biol. 23, 207–236. https://doi.org/10.1146/annurev.cellbio.22.011105.102609

Jullien, P.E., Kinoshita, T., Ohad, N., Berger, F., 2006. Maintenance of DNA Methylation during the Arabidopsis Life Cycle Is Essential for Parental Imprinting. Plant Cell 18, 1360–1372. https://doi.org/10.1105/tpc.106.041178

Karimi, M., Inzé, D., Depicker, A., 2002. GATEWAY™ vectors for Agrobacterium-mediated plant transformation. Trends Plant Sci. 7, 193–195. https://doi.org/10.1016/S1360-1385(02)02251-3

Kinoshita, T., Harada, J.J., Goldberg, R.B., Fischer, R.L., 2001. Polycomb repression of flowering during early plant development. Proc. Natl. Acad. Sci. 98, 14156–14161. https://doi.org/10.1073/pnas.241507798

Kinoshita, T., Yadegari, R., Harada, J.J., Goldberg, R.B., Fischer, R.L., 1999. Imprinting of the MEDEA Polycomb Gene in the Arabidopsis Endosperm. Plant Cell 11, 1945–1952. https://doi.org/10.1105/tpc.11.10.1945

Kirmizis, A., Bartley, S.M., Kuzmichev, A., Margueron, R., Reinberg, D., Green, R., Farnham, P.J., 2004. Silencing of human polycomb target genes is associated with methylation of histone H3 Lys 27. Genes Dev. 18, 1592–1605. https://doi.org/10.1101/gad.1200204

Kiyosue, T., Ohad, N., Yadegari, R., Hannon, M., Dinneny, J., Wells, D., Katz, A., Margossian, L., Harada, J.J., Goldberg, R.B., Fischer, R.L., 1999. Control of fertilization-independent endosperm development by the MEDEA polycomb gene in Arabidopsis. Proc. Natl. Acad. Sci. 96, 4186–4191. https://doi.org/10.1073/pnas.96.7.4186

Köhler, C., Hennig, L., Bouveret, R., Gheyselinck, J., Grossniklaus, U., Gruissem, W., 2003a. Arabidopsis MSI1 is a component of the MEA/FIE Polycomb group complex and required for seed development. EMBO J. 22, 4804–4814. https://doi.org/10.1093/emboj/cdg444

Köhler, C., Hennig, L., Spillane, C., Pien, S., Gruissem, W., Grossniklaus, U., 2003b. The Polycomb-group protein MEDEA regulates seed development by controlling expression of the MADS-box gene PHERES1. Genes Dev. 17, 1540–1553. https://doi.org/10.1101/gad.257403

Lau, M.M., Stewart, C.E., Liu, Z., Bhatt, H., Rotwein, P., Stewart, C.L., 1994. Loss of the imprinted IGF2/cation-independent mannose 6-phosphate receptor results in fetal overgrowth and perinatal lethality. Genes Dev. 8, 2953–2963. https://doi.org/10.1101/gad.8.24.2953

Laugesen, A., Højfeldt, J.W., Helin, K., 2016. Role of the Polycomb Repressive Complex 2 (PRC2) in Transcriptional Regulation and Cancer. Cold Spring Harb. Perspect. Med. 6. https://doi.org/10.1101/cshperspect.a026575

Leighton, P.A., Ingram, R.S., Eggenschwiler, J., Efstratiadis, A., Tilghman, S.M., 1995. Disruption of imprinting caused by deletion of the H19 gene region in mice. Nature 375, 34–39. https://doi.org/10.1038/375034a0

Leroy, O., Hennig, L., Breuninger, H., Laux, T., Köhler, C., 2007. Polycomb group proteins function in the female gametophyte to determine seed development in plants. Development 134, 3639–3648. https://doi.org/10.1242/dev.009027

Liao, C.-Y., Smet, W., Brunoud, G., Yoshida, S., Vernoux, T., Weijers, D., 2015. Reporters for sensitive and quantitative measurement of auxin response. Nat. Methods 12, 207–210. https://doi.org/10.1038/nmeth.3279

Lodha, M., Marco, C.F., Timmermans, M.C.P., 2013. The ASYMMETRIC LEAVES complex maintains repression of KNOX homeobox genes via direct recruitment of Polycomb-repressive complex2. Genes Dev. 27, 596–601. https://doi.org/10.1101/gad.211425.112

Loubiere, V., Martinez, A.-M., Cavalli, G., 2019. Cell Fate and Developmental Regulation Dynamics by Polycomb Proteins and 3D Genome Architecture. BioEssays 41, 1800222. https://doi.org/10.1002/bies.201800222

Margueron, R., Reinberg, D., 2011. The Polycomb Complex PRC2 and its Mark in Life. Nature 469, 343–349. https://doi.org/10.1038/nature09784

Maruyama, D., Hamamura, Y., Takeuchi, H., Susaki, D., Nishimaki, M., Kurihara, D., Kasahara, R.D., Higashiyama, T., 2013. Independent Control by Each Female Gamete Prevents the Attraction of Multiple Pollen Tubes. Dev. Cell 25, 317–323. https://doi.org/10.1016/j.devcel.2013.03.013

Meijer, M., Murray, J.A.H., 2000. The role and regulation of D-type cyclins in the plant cell cycle. Plant Mol. Biol. 43, 621–633. https://doi.org/10.1023/A:1006482115915

Möller, B., Weijers, D., 2009. Auxin Control of Embryo Patterning. Cold Spring Harb. Perspect. Biol. 1. https://doi.org/10.1101/cshperspect.a001545

O’Carroll, D., Erhardt, S., Pagani, M., Barton, S.C., Surani, M.A., Jenuwein, T., 2001. The Polycomb-Group GeneEzh2 Is Required for Early Mouse Development. Mol. Cell. Biol. 21, 4330–4336. https://doi.org/10.1128/MCB.21.13.4330-4336.2001

O’Dor, E., Beck, S.A., Brock, H.W., 2006. Polycomb group mutants exhibit mitotic defects in syncytial cell cycles of Drosophila embryos. Dev. Biol. 290, 312–322. https://doi.org/10.1016/j.ydbio.2005.11.015

Ohad, N., Yadegari, R., Margossian, L., Hannon, M., Michaeli, D., Harada, J.J., Goldberg, R.B., Fischer, R.L., 1999. Mutations in FIE, a WD Polycomb Group Gene, Allow Endosperm Development without Fertilization. Plant Cell 11, 407–415. https://doi.org/10.1105/tpc.11.3.407

Oktaba, K., Gutiérrez, L., Gagneur, J., Girardot, C., Sengupta, A.K., Furlong, E.E.M., Müller, J., 2008. Dynamic Regulation by Polycomb Group Protein Complexes Controls Pattern Formation and the Cell Cycle in Drosophila. Dev. Cell 15, 877–889. https://doi.org/10.1016/j.devcel.2008.10.005

Palovaara, J., Saiga, S., Wendrich, J.R., van ‘t Wout Hofland, N., van Schayck, J.P., Hater, F., Mutte, S., Sjollema, J., Boekschoten, M., Hooiveld, G.J., Weijers, D., 2017. Transcriptome dynamics revealed by a gene expression atlas of the early Arabidopsis embryo. Nat. Plants 3, 894–904. https://doi.org/10.1038/s41477-017-0035-3

Pasini, D., Bracken, A.P., Jensen, M.R., Denchi, E.L., Helin, K., 2004. Suz12 is essential for mouse development and for EZH2 histone methyltransferase activity. EMBO J. 23, 4061–4071. https://doi.org/10.1038/sj.emboj.7600402

Pires, N.D., Bemer, M., Müller, L.M., Baroux, C., Spillane, C., Grossniklaus, U., 2016. Quantitative Genetics Identifies Cryptic Genetic Variation Involved in the Paternal Regulation of Seed Development. PLOS Genet. 12, e1005806. https://doi.org/10.1371/journal.pgen.1005806

Pires, N.D., Grossniklaus, U., 2014. Different yet similar: evolution of imprinting in flowering plants and mammals. F1000Prime Rep. 6. https://doi.org/10.12703/P6-63

Raissig, M.T., Bemer, M., Baroux, C., Grossniklaus, U., 2013. Genomic Imprinting in the Arabidopsis Embryo Is Partly Regulated by PRC2. PLoS Genet. 9. https://doi.org/10.1371/journal.pgen.1003862

Ron, M., Alandete Saez, M., Eshed Williams, L., Fletcher, J.C., McCormick, S., 2010. Proper regulation of a sperm-specific cis-nat-siRNA is essential for double fertilization in Arabidopsis. Genes Dev. 24, 1010–1021. https://doi.org/10.1101/gad.1882810

Sarkar, A.K., Luijten, M., Miyashima, S., Lenhard, M., Hashimoto, T., Nakajima, K., Scheres, B., Heidstra, R., Laux, T., 2007. Conserved factors regulate signalling in Arabidopsis thaliana shoot and root stem cell organizers. Nature 446, 811–814. https://doi.org/10.1038/nature05703

Schlereth, A., Möller, B., Liu, W., Kientz, M., Flipse, J., Rademacher, E.H., Schmid, M., Jürgens, G., Weijers, D., 2010. MONOPTEROS controls embryonic root initiation by regulating a mobile transcription factor. Nature 464, 913–916. https://doi.org/10.1038/nature08836

Scott, R.J., Vinkenoog, R., Spielman, M., Dickinson, H.G., 1998. Medea: murder or mistrial? Trends Plant Sci. 3, 460–461. https://doi.org/10.1016/S1360-1385(98)01351-X

Simonini, S., Deb, J., Moubayidin, L., Stephenson, P., Valluru, M., Freire-Rios, A, Sorefan, K., Weijers, D., Friml, J., Østergaard, L., 2016. A noncanonical auxin-sensing mechanism is required for organ morphogenesis in Arabidopsis. Genes Dev. 30, 2286–2296. https://doi.org/10.1101/gad.285361.116

Skene, P.J., Henikoff, J.G., Henikoff, S., 2018. Targeted in situ genome-wide profiling with high efficiency for low cell numbers. Nat. Protoc. 13, 1006–1019. https://doi.org/10.1038/nprot.2018.015

Sozzani, R., Cui, H., Moreno-Risueno, M.A., Busch, W., Van Norman, J.M., Vernoux, T., Brady, S.M., Dewitte, W., Murray, J. a. H., Benfey, P.N., 2010. Spatiotemporal regulation of cell-cycle genes by SHORTROOT links patterning and growth. Nature 466, 128–132. https://doi.org/10.1038/nature09143

Spillane, C., Schmid, K.J., Laoueillé-Duprat, S., Pien, S., Escobar-Restrepo, J.-M., Baroux, C., Gagliardini, V., Page, D.R., Wolfe, K.H., Grossniklaus, U., 2007. Positive darwinian selection at the imprinted MEDEA locus in plants. Nature 448, 349–352. https://doi.org/10.1038/nature05984

Stent, G.S., 1985. The Role of Cell Lineage in Development. Philos. Trans. R. Soc. Lond. B. Biol. Sci. 312, 3–19.

Truernit, E., Bauby, H., Dubreucq, B., Grandjean, O., Runions, J., Barthélémy, J., Palauqui, J.-C., 2008. High-Resolution Whole-Mount Imaging of Three-Dimensional Tissue Organization and Gene Expression Enables the Study of Phloem Development and Structure in Arabidopsis. Plant Cell 20, 1494–1503. https://doi.org/10.1105/tpc.107.056069

Vielle-Calzada, J.-P., Thomas, J., Spillane, C., Coluccio, A., Hoeppner, M.A., Grossniklaus, U., 1999. Maintenance of genomic imprinting at the Arabidopsis medea locus requires zygotic DDM1 activity. Genes Dev. 13, 2971–2982.

Vieten, A., Vanneste, S., Wisniewska, J., Benková, E., Benjamins, R., Beeckman, T., Luschnig, C., Friml, J., 2005. Functional redundancy of PIN proteins is accompanied by auxin-dependent cross-regulation of PIN expression. Dev. Camb. Engl. 132, 4521–4531. https://doi.org/10.1242/dev.02027

Wang, L., Balas, B., Christ-Roberts, C.Y., Kim, R.Y., Ramos, F.J., Kikani, C.K., Li C., Deng, C., Reyna, S., Musi, N., Dong, L.Q., DeFronzo, R.A., Liu, F., 2007. Peripheral Disruption of the Grb10 Gene Enhances Insulin Signaling and Sensitivity In Vivo. Mol. Cell. Biol. 27, 6497–6505. https://doi.org/10.1128/MCB.00679-07

Weijers, D., Dijk, M.F., Vencken, R.-J., Quint, A., Hooykaas, P., Offringa, R., 2001. An Arabidopsis Minute-like phenotype caused by a semi-dominant mutation in a RIBOSOMAL PROTEIN S5 gene. Development 128, 4289–4299.

Wysocka-Diller, J.W., Helariutta, Y., Fukaki, H., Malamy, J.E., Benfey, P.N., 2000. Molecular analysis of SCARECROW function reveals a radial patterning mechanism common to root and shoot. Development 127, 595–603.

Yan, H., Chen, D., Wang, Y., Sun, Y., Zhao, J., Sun, M., Peng, X., 2016. Ribosomal protein L18aB is required for both male gametophyte function and embryo development in Arabidopsis. Sci. Rep. 6, 1–12. https://doi.org/10.1038/srep31195

Zheng, X., Gehring, M., 2019. Low-input chromatin profiling in Arabidopsis endosperm using CUT&RUN. Plant Reprod. 32, 63–75. https://doi.org/10.1007/s00497-018-00358-1

